# Predation by protists influences the temperature response of microbial communities

**DOI:** 10.1101/2021.04.08.439073

**Authors:** Jennifer D. Rocca, Andrea Yammine, Marie Simonin, Jean P. Gibert

## Abstract

Temperature strongly influences microbial community structure and function, which in turn contributes to the global carbon cycle that can fuel further warming. Recent studies suggest that biotic interactions amongst microbes may play an important role in determining the temperature responses of these communities. However, how microbial predation regulates these communities under future climates is still poorly understood. Here we assess whether predation by one of the most important bacterial consumers globally – protists – influences the temperature response of a freshwater microbial community structure and function. To do so, we exposed these microbial communities to two cosmopolitan species of protists at two different temperatures, in a month-long microcosm experiment. While microbial biomass and respiration increased with temperature due to shifts in microbial community structure, these responses changed over time and in the presence of protist predators. Protists influenced microbial biomass and function through effects on community structure, and predation actually reduced microbial respiration rate at elevated temperature. Indicator species and threshold indicator taxa analyses showed that these predation effects were mostly determined by phylum-specific bacterial responses to protist density and cell size. Our study supports previous findings that temperature is an important driver of microbial communities, but also demonstrates that predation can mediate these responses to warming, with important consequences for the global carbon cycle and future warming.

## INTRODUCTION

Understanding the biotic factors that influence global climate change is one of the most pressing goals of ecology (Van der Putten et al., 2010). Doing so hinges on better understanding the biotic and abiotic feedbacks that determine global carbon cycling (Jackson et al., 2017). While several of the biotic mechanisms that lead to global releases and sequestration of carbon are well documented (Nielsen et al., 2014; van den Hoogen et al., 2020), only a subset of those are currently taken into account in predictive Earth System Models (Sulman et al., 2019).

Microbial organisms comprise 14% of all existing biomass on Earth (Bar-On et al., 2018; Flemming & Wuertz, 2019), while the entirety of the Animal Kingdom, for comparison, only represents 0.3% (Bar-On et al., 2018). Microbial decomposition is responsible for the recycling of organic matter back into food webs, thus partly subsidizing the flux of energy and matter in all ecosystems (Cordone et al., 2020; Mougi, 2020). Through respiration and decomposition of existing carbon pools, microorganisms are largely regarded as one of the most important biotic controls of the global carbon cycle (Bardgett et al., 2008; Gougoulias et al., 2014; Jansson & Hofmockel, 2020; Schimel & Schaeffer, 2012; Wang, 2018). Additionally, respiration rates are well-known to increase with temperature among ectotherms and unicellular organisms (Bond-Lamberty et al., 2018; DeLong et al., 2017; Gillooly et al., 2001), potentially resulting in a scenario where warming begets more warming. However, temperature-mediated increases in respiration rates often plateau –or even decline– over time in microbial communities (Bradford, 2013; Crowther & Bradford, 2013; Ye et al., 2020; Yergeau et al., 2012), although not always (Hartley et al., 2008; Zimmermann et al., 2012). These diverse responses thus show there is much that still needs to be understood about the processes regulating microbial respiration and function in the wild.

Temperature often mediates ecological interactions (Bernhardt et al., 2018; Binzer et al., 2012; Dell et al., 2014; Garzke et al., 2019; Uszko et al., 2017). The strength of feeding interactions, in particular, increases with temperature, as feeding rates increase among consumers to compensate for increasing metabolic demands (Dell et al., 2011; Gillooly et al., 2001). Stronger predation in turn leads to declines in prey abundance and total biomass (Barneche et al., 2021; DeLong & Lyon, 2020; Garzke et al., 2019; Gilbert et al., 2014). Because gross respiration rates are determined by standing biomass, temperature effects on predation may ultimately influence ecosystem-level processes such as community respiration rates (O’Connor et al., 2009), and thus mediate the temperature response of microbial respiration rates worldwide. In particular, bacterivory is a dominant factor leading to microbial biomass loss (Baltar et al., 2016; Pedrós-Alió et al., 2000; Rgens & Massana, 2008), which has been proposed to influence soil respiration rates (Gao et al., 2019), and shown to affect litter decomposition in soils across temperatures (Geisen et al., 2021; Sulman et al., 2018).

Recent efforts have mapped the global distribution of nematodes – a major group of microbial predators (Nielsen et al., 2014; van den Hoogen et al., 2019, 2020). These global maps represent a step forward in clarifying the role of feeding interactions in the temperature responses of microbial communities. However, with a global biomass 200 times larger than that of nematodes (Bar-On et al., 2018), unicellular eukaryotes - collectively known as ‘protists’ - likely play a major role in regulating microbial communities at global scales (Oliverio et al., 2020) through bacterivory (Erktan et al., 2020; Gao et al., 2019). Ciliate protists, in particular, are well-known bacterivores (Foissner & Berger, 1996), their population dynamics and feeding interactions are strongly temperature-dependent (DeLong & Lyon, 2020), and they are present in all major ecosystems (Foissner & Berger, 1996; Oliverio et al., 2020). As such, predation of microorganisms by protists can mediate the temperature response of microbial communities (Gao et al., 2019; Geisen et al., 2021), although this phenomenon has only been shown for one species of protist in soils (Geisen et al., 2021). If general, this process has the potential to strongly influence microbial respiration worldwide under warmer temperatures (Crowther et al., 2015; Geisen et al., 2018). Additionally, microbial predators respond to environmental conditions themselves in multiple ways, including changes in feeding rates (DeLong & Lyon, 2020; Englund et al., 2011) and changes in the traits that influence predation (Atkinson et al., 2003; Dell et al., 2011; Gibert et al., 2016), which add complexity to an already complex problem.

Here, we examine the potential interactive effects of protist predation and temperature on microbial diversity, biomass, and total respiration rates, by incubating a microbial community from a local ephemeral pond in the presence and absence of two cosmopolitan ciliate protists of different size and at different temperatures. We ask: 1) how does temperature influence microbial community biomass, structure, and function?, 2) how does protist presence influence the temperature response of microbial community biomass, structure, and function?, and 3) are there direct temperature responses of the protists influencing their effects on the microbial community they feed upon?

We hypothesize that microbial biomass and function should increase with temperature, although we expect that effect to plateau over time (Bradford, 2013; Crowther & Bradford, 2013; Ye et al., 2020; Yergeau et al., 2012). We expect protist predation to decrease overall microbial biomass (Glücksman et al., 2010; Hahn & Höfle, 2001), which could reduce total respiration rates. We also hypothesize that predation effects should be dependent on protist size, as feeding rates are well-known to increase with body size (Zaoli et al., 2019). While little is known about the exact diet of these protists, we expect them to differ, at least minimally because of gape-limitation (Glücksman et al., 2010): larger protists should be able to consume the same species that smaller protists do, plus some biofilm or colony-making microbial taxa. Consequently, we hypothesize that differential consumption of microbial species by different protist species should result in changes in microbial composition (Glücksman et al., 2010; Hahn & Höfle, 2001), that may lead to changes in community structure and function. Last, we predict that protists themselves may respond to the imposed treatments by decreasing body size with temperature (temperature-size rule; (D. Atkinson, 1994; Atkinson et al., 2006). These changes may in turn influence how they interact with the microbial community.

## METHODS

### Water collection, microcosm setup and incubation

We collected 40 L of surface water from an ephemeral freshwater pond at Duke Forest (Gate 9, 36.019139, −78.987698, Durham, NC). To isolate the aquatic microbial community used in our experiment, we filtered the entire water sample through autoclaved filters (11μm pore size, Whatman) to remove debris, metazoans, and larger protists; then we filtered through sterile GF/A filters (1.6μm pore size, Whatman) to remove flagellates and other smaller organisms. Smaller microbes were retained in the filtrate. Removal of protists and metazoans from the source pond microbial community was confirmed by visual inspection using a stereomicroscope (Leica M205C), then re-confirmed on control microcosms at the end of the experiment using fluid imaging (detailed below). The microbial communities were incubated in 250mL acid-washed and autoclaved borosilicate jars filled with a mixture of pond water filtrate (2/3, or 133mL), and Carolina Biological Protist culture medium (1/3, or 67mL) plus two wheat seeds to serve as a carbon source ((Altermatt et al., 2015), for a total of 140 replicate microcosms. We included a negative control (n=1), containing the same volume of sterile protist media, to confirm axenic conditions throughout the incubation and subsequent processing. All microcosms were incubated in Percival AL-22L2 growth chambers (Percival Scientific, Perry, Iowa) at 22°C, 10% light intensity (1700 lux), 75% humidity, and a 16:8 hr day-night cycle (day length at time of collection). After a seven day pre-incubation period, to allow equilibration of the microbial communities, we harvested 20 microcosms as positive controls to assess the effects of incubation on microbial composition, relative to the original pond microbial community. The negative control was extracted for genomic DNA alongside these 20 samples (*detailed below*).

### Experimental treatments

After the initial seven-day incubation period, microcosms were randomly assigned to treatments in a fully factorial experimental design with two levels of temperature (22°C, i.e., the water temperature on the day of collection; or 25°C, a warming scenario simulating a +3°C increase in temperature), and three levels of protist predation (no protist, presence of *Tetrahymena pyriformis*, or presence of *Colpidium sp.*). We used *Tetrahymena pyriformis* (hereafter *Tetrahymena*) and *Colpidium sp.* (hereafter *Colpidium*), due to their putative generalist bacterivore habits (Foissner & Berger, 1996), their cosmopolitan distribution (Elliott, 1970), and, hence, their likelihood of playing a pivotal role in mediating the temperature response of microbial communities worldwide. Also, these protists have a large size difference (20-70 μm for *Tetrahymena* vs 60-120 μm of *Colpidium*) which theory predicts should lead to differences in feeding and interaction strengths with their bacterial prey (Englund et al., 2011; Rall et al., 2012; Zaoli et al., 2019).

Protists were introduced by pipetting 0.5mL of well-mixed protist stock cultures into experimental microcosms. To control for the introduction of the microbes already occurring in the protist cultures, we also added the same volume of a homogenized protist stock media, filtered of *Tetrahymena* and *Colpidium* cells, into all microcosms. The microcosms were thus assigned to one of 6 possible treatments: 1) 22°C, no protists; 2) 25°C, no protists; 3) 22°C, *Tetrahymena*; 4) 25°C, *Tetrahymena*; 5) 22°C, *Colpidium*; 6) 25°C, *Colpidium*. Half of the microcosms in each treatment were harvested at Day 12 and the remaining half at Day 24, to assess whether observed responses changed over time in systematic ways.

### Microbial Biomass and Community Respiration

We quantified total microbial biomass, microbial diversity, and total community respiration to assess the joint impacts of temperature, time, and protist predation on microbial community structure and function. As a proxy for total biomass, we measured the optical density at 600nm wavelength (or OD600; (Beal et al., 2020) of each microcosm (1/3 dilutions), using a Jenway 3505 UV Spectrophotometer (Cole-Parmer, Vernon-Hills, IL, USA). Larger OD600 values (higher absorbance+scattering) indicate higher total microbial biomass (Beal et al., 2020).

We determined total community respiration using an optode-based real time OXY-4 SMA respirometer (PreSens, Regensburg, Germany; (Altermatt et al., 2015; DeLong & Vasseur, 2012). Respiration rate was measured on 22 mL subsamples for 30 minutes, after a 30 minute acclimation period, on a subset of all microcosms (n=72). This was done at their original experimental temperature and the cross-treatment temperature (n=36 microcosms, with two measurements each) to disentangle long term temperature effects from short-term impacts. Respiration rates were estimated as the rate of change (slope) of the estimated oxygen concentration over time (in μmol O2/min, Figures S1, S2, Appendix 1). Respiration rates did not differ significantly between the two temperatures at which they were measured (effect = − 0.01±0.12SE, *p*=0.96), so readings for both temperatures from a single microcosm were averaged for subsequent analyses. To assess whether total biomass or community respiration changed with experimental treatment, we used linear models with protist presence (no protist, *Tetrahymena* or *Colpidium*), time (12 or 24 days), and temperature (22 or 25°C) as explanatory variables (and their possible interactions), and either biomass or respiration rates as response variables in R v4.0.2; R Core Team, 2013). All measures of biomass, function and microbial community structure were measured at days 12 and 24.

### Protist abundances and traits

To disentangle potential effects of protist presence and abundance on microbial communities, we estimated protist population sizes through fluid imaging of 3 mL subsamples out of four microcosms from each treatment using a FlowCam (Fluid Imaging Technologies, Scarborough, ME, USA). Fluid imaging also yields high-resolution measurements of multiple cell traits like cellular volume and shape, and optical properties of the cells (Gibert et al., 2017). We used ensuing trait data to assess potential responses of the protists to imposed experimental conditions as well as potential responses of the microbial communities to both protist traits and densities (detailed below). We focused on nine different phenotypic characteristics: five measurements of shape and size (length, area, volume, circularity, and aspect ratio) and four measurements of optical properties of the cells (sigma-intensity and three components of hue: Red/Green, Red/Blue and Blue/Green ratios). These measurements were taken on days 12, and 24. We assessed whether protist population counts changed with either temperature or time using linear models, and we assessed whether protist phenotypes responded over time and with temperature using perMANOVA (Anderson, 2001, 2017) using the *vegan* package in R (v2.5.6; (Oksanen et al., 2019).

### Amplicon Sequence Data Processing and Microbial Community Structure Analysis

We used 16S rRNA amplicon sequencing to examine the impacts of temperature, time, and protists, on microbial community structure. After the incubation period (12 or 24 days), we collected the microbial communities by filtering 200mL from each microcosm onto gamma-irradiated 0.2μm nitrocellulose membranes (Advantec, Taipei, Taiwan) and stored the filters at - 20°C until DNA extraction. Total genomic DNA was extracted from each filter using DNeasy PowerWater DNA Extraction Kits (Qiagen, Hilden, Germany), modified with a heating step (60°C) before the initial vortexing step to maximize lysis across different microbial cell types. We fluorometrically quantified the genomic DNA concentrations with Qubit (Thermo Fisher, Waltham, MA, USA), and sent an equimolar set of genomic DNA samples to the Research Technology Support Facility (RTSF) at Michigan State University for amplicon prep and sequencing. We targeted the V4 hypervariable region of the 16S rRNA gene using the standard 515F/806R universal primers with 12 bp Golay barcodes (Caporaso et al., 2011). RTSF sequenced our samples with Illumina MiSeq (PE 250 bp, V2 chemistry), and returned 9,069,268 raw reads (average/sample: 62,981 reads), publicly available at EMBL-ENA under project accession number: PRJEB44142 (ERS6200133 - ERS6200276).

We processed the raw fastq sequence data through Dada2 (v1.16.0; (Callahan et al., 2016) in R (v4.0.2; R Core Team, 2013), to trim and filter low quality sequence reads and calculate error rates for denoising and merging the pair-ends into 3801 non-chimeric representative amplicon sequence variants, or ASVs. These representative ASVs were further curated with Lulu to reduce artificially inflated diversity due to amplification and sequencing errors, resulting in 1423 representative ASVs (Frøslev et al., 2017).

We taxonomically identified chloroplast and mitochondrial 16S rRNA sequences using the Silva 138 reference taxonomy (Quast et al., 2013), removing 315 ASVs for a final 16S rRNA representative set of 1108 ASVs. This final representative ASV set was aligned to the Silva 138 NR full-length 16S rRNA alignment with MAFFT (Katoh & Frith, 2012) using default settings, and subsequently trimmed and masked to the V4 region. We then updated the trimmed V4 alignment to estimate a phylogeny using the iterative algorithm of PASTA (Mirarab et al., 2015), and used this final set of ASVs to update the corresponding sample ASV community table.

For alpha diversity estimates, we rarefied all microbial 16S rRNA samples to a sequencing depth of 8600 to maximize sampling depth while retaining the majority of samples. We treated the data as compositional for all other analyses by using a variance-stabilizing transformation (VST) of the ASV community table without singleton ASVs (Gloor et al., 2017), using DESeq2 (v1.12.3; (McMurdie & Holmes, 2013). We calculated a comprehensive range of alpha-diversity indices, from observed ASV richness and Shannon-Weiner to Pielou’s evenness index and abundance-weighted phylogenetic diversity (weighted Unifrac) on each rarefied sample using ‘core-metrics-phylogenetic’ in Qiime2 (v2020.8; (Bolyen et al., 2019). We used ANOVA to test for individual and interactive effects of experimental treatments on alpha diversity of the microbial communities.

To examine the impacts of temperature, incubation time, and protists on the overall structure of the microbial communities, we used principal component analysis on an abundance-weighted bray-curtis distance matrix of the VST table in *vegan* R (v2.5-6; (Oksanen et al., 2019). We tested for individual and interactive treatment effects with a perMANOVA using the adonis() function in *vegan* R (v2.5-6; (Oksanen et al., 2019). We performed multilevel pairwise comparisons of the community data using the pairwise.adonis() function in the pairwiseAdonis R package (v0.0.1; (Martinez Arbizu, 2017). We also examined differences among group community variation –or spread in ordination space– using the betadisper() function (PERMDISP2; (Anderson et al., 2006) in *vegan* R (v2.5-6; (Oksanen et al., 2019), which uses multivariate homogeneity of group dispersions, or the multivariate form of a Levene’s test (Anderson, 2006; O’Neill & Mathews, 2000).

For treatments imposing significant changes in microbial community structure, we also identified potential positive or negatively responding microbes (ASVs). We employed two distinctive methods of indicator taxa analysis: categorical *indicspecies* in R (De Cáceres & Legendre, 2009) and a direct gradient analysis, threshold indicator taxa analysis (TITAN) in R (Baker & King, 2010). With indicspecies, we identified ASVs with significant differential abundance patterns by treatment level using multi-level pattern analysis with the multipatt() function (De Cáceres & Legendre, 2009). In contrast, with TITAN, we identified microbial ASVs significantly associated with changes in measured protist phenotypic traits, specifically protist size, and protist abundance, by separately regressing each microbial ASV against each protist trait. TITAN outputs provided pure and reliable positive and negative indicator ASVs (75% purity and reliability thresholds), as well as individual and community-level abundance thresholds, for each protist trait gradient. All data and code used for these analyses, can be accessed at https://github.com/JPGibert/Microbial_Responses_Prots_Temp, unless otherwise stated.

## RESULTS

### Temperature and time affect microbial biomass, function, and structure

Temperature and incubation time had interactive effects on total microbial biomass: warmer temperature resulted in larger OD600, but that effect disappeared over time (Fig 1a, effect_Temp_ = 0.08±0.04SE, *p*=0.08; effect_Time_ = 0.10±0.04SE, *p*=0.019; effect_Temp x Time_ = –0.14±0.06SE, *p*=0.02, more details in Appendix 2). Respiration rate also showed significant interactive effects of time and temperature similar to those found for bacterial biomass: respiration increased with temperature at first, but that effect was reversed over time (Fig 1b, effect_Temp_ = 0.62±0.29SE, *p*=0.036; effect_Time_ = 0.46±0.20SE, *p*=0.025; effect_Temp x Time_ = –0.90±0.29SE, *p*=0.003, Appendix 2).

**Figure 1.**
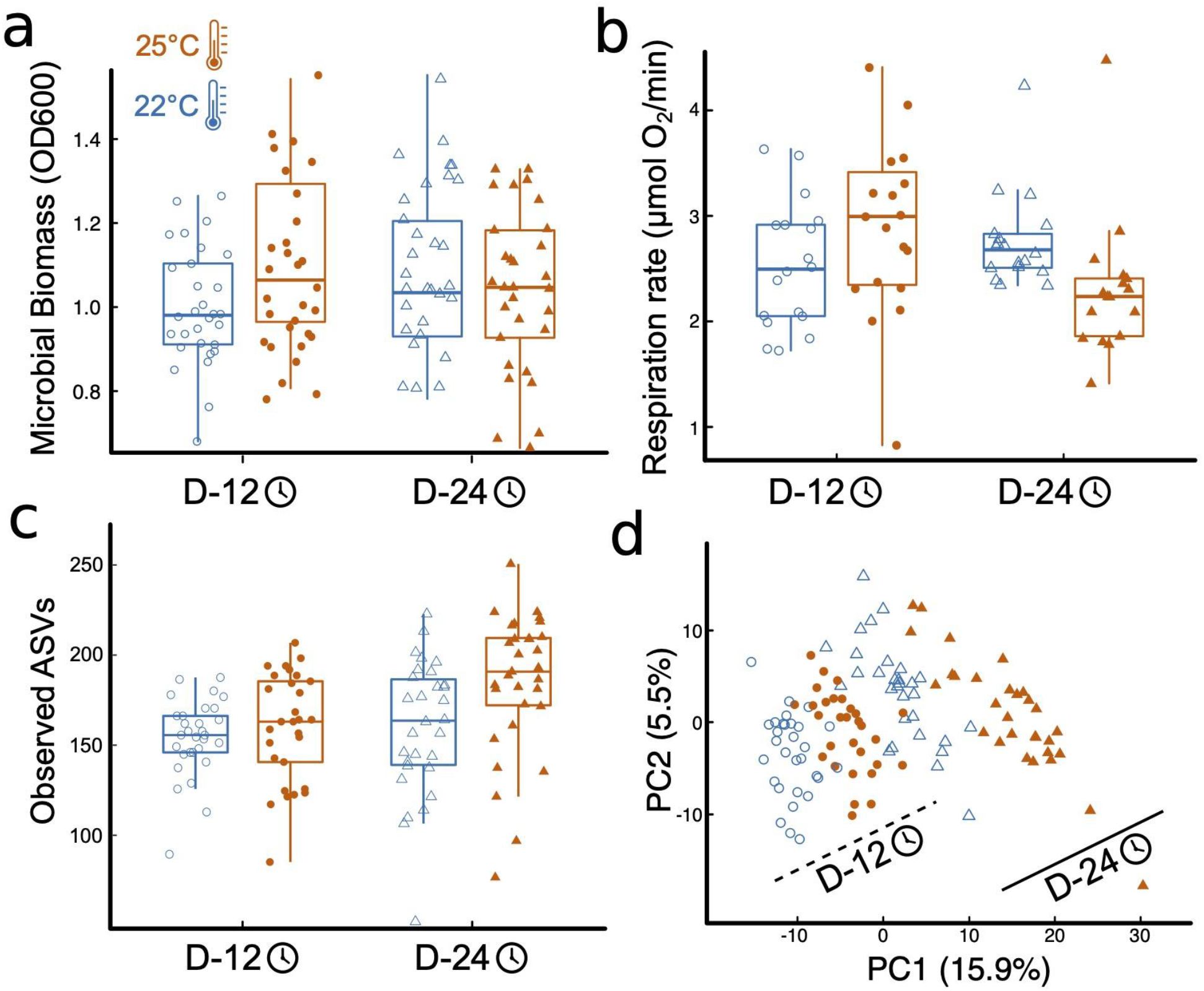
Temperature impacts on microbial function and community structure. (a) Boxplots of the treatment effects on total microbial biomass, (b) the effects on total microbial respiration, here as oxygen consumption, (c) the effects on microbial community richness, here observed ASVs, and (d) the effects of temperature and time on the microbial community structure. Shapes represent incubation time: Day12 - circles, Day 24 - triangles; and temperature marked by point fill: 22°C - empty point, 25°C - color filled point.

Temperature and time, but not their interaction, influenced microbial community structure, with increased alpha- and beta-diversity with elevated temperature and with time. (Fig 1c, Appendix 3). Mimicking biomass and respiration results, microbial community structure was significantly affected by temperature, time, and their interaction (Fig 1d). The microbial communities were primarily structured by incubation time, with 13.3% of the variation explained by community shifts from Day 12 to Day 24 harvest (*p=.001*). Temperature (22°C vs. 25°C) explained 5.5% of the community variation across all harvesting time points (*p=.001*, Fig 1d), while the interaction of temperature and time explained 3.2% of the variation in microbial community structure (*p=.001*, Fig 1d). Post-hoc tests revealed that all four treatment combinations resulted in significantly distinct groupings of microbial community structure (*p.adj=.006*). Finally, increased temperature and incubation time imposed significant increases to beta-dispersion (temp: *p.adj=.003*, time: *p.adj=.001*), with wider spread in group dispersion at the warmer temperature in the early harvested microbiomes (*p.adj=.045*) (Fig 1d).

### Effects of protist predation on the microbial function and community structure

The larger protist (*Colpidium*) significantly reduced OD600 biomass relative to the no protist treatment, while the smaller *Tetrahymena* did not (Fig 2a, effect_Colp_= −0.09±0.04SE, *p*=0.015; effect_Tetra_= −0.05±0.04SE, *p*=0.19). Protist effects on OD600 biomass did not interact with time or temperature (Table S1, Fig S3, Appendix 2). *Tetrahymena* had no noticeable effect on microbial respiration rate either (Fig 2b, effect_Tetra_= −0.05±0.04SE, *p*=0.19). However, *Colpidium* had a significant effect on respiration that interacted with temperature: in the presence of *Colpidium,* total respiration was lower at 25°C than at 22°C (effect_Colp_= 0.99±0.25SE, *p*<0.01; effect_Colp x Temp_= −0.87±0.36SE, *p*=0.017; Fig 2b).

**Figure 2.**
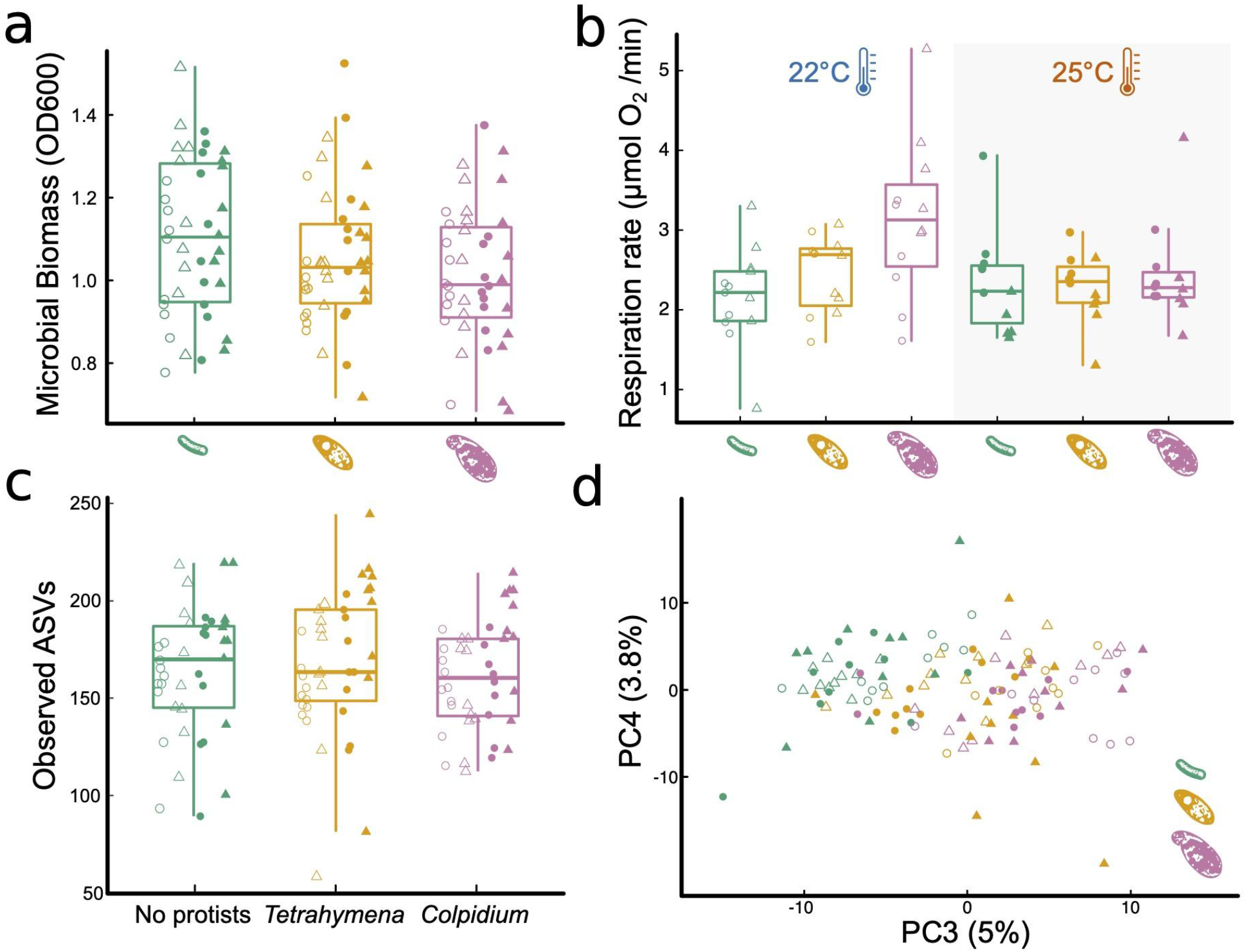
Influence of protist presence and species on pond microbial community function and community structure. The effect of the absence of protists (green), *Tetrahymena* (yellow) or *Colpidium* (fuchsia) on: (a) optical density (600nm) as a proxy for total microbial biomass, (b) total microbial respiration (O2 consumption rate), (c) microbial community richness (observed ASVs), and (d) a principal component analysis of microbial community structure. Shapes represent incubation time: Day12 - circles, Day 24 - triangles; and temperature marked by point fill: 22°C - empty point, 25°C - color filled point.

Predator presence had no significant effects on any of the measured alpha diversity indices of the microbial communities (ASV richness, *p = 0.6*; phylogenetic diversity, *p = 0.8*; Shannon-Wiener diversity, *p = 0.5*; or community evenness, *p = 0.4*, Fig 2c, Appendix 3). However, both protists significantly affected microbial community composition (Fig 2d), although neither one interacted with temperature or time (protist and temperature interaction: p=0.22; protist and time interaction: p=0.25; protist, time and temperature interaction: p=0.60). Protist treatments explained 6.7% of the variation in microbial community structure (*p=.001*). Microbial communities exposed to either protist species significantly differed from the no protist microbial communities (*Tetrahymena*: *p.adj=.001*; *Colpidium*: *p.adj=.001*), and also differed between the two protist species treatments (*p.adj=.004*). Group dispersion analysis of beta diversity revealed no differences in the degree of group variation in community structure among protist treatment levels (*p.adj=.239*).

### Feedbacks on protist abundance and traits

Protist abundance increased over time (Fig 3a effect_Colp_ = 172.05±46.77SE, *p*=0.002; effect_Tetra_ = 344.9±132.9SE, *p*=0.02), as expected, but final abundance was independent from temperature (Fig 3b; effect_Colp_ = −16.43±66.66SE, *p*=0.89; effect_Tetra_ = 120.9±160.3SE, *p*=0.46). This indicates that temperature effects on protist abundances are unlikely to explain, alone, observed effects on microbial communities and respiration rate. Both time and temperature influenced protist traits independently and interactively (Fig 3c-f, Tables S4-S7, Appendix 4), but each species responded in slightly different ways. Contrary to the temperature-size rule, *Colpidium sp.* responded to increasing temperatures by becoming larger and more elongated (Fig 3c, perMANOVA *p*<0.01, Table S4, S5, Appendix 4). On the other hand, *Tetrahymena* response was consistent with a temperature-size rule, becoming smaller and rounder with temperature (Fig 3e, perMANOVA *p*<0.01,Table S6, S7, Appendix 4). Over time, however, *Colpidium* got smaller and shorter (Fig 3d, perMANOVA *p*<0.01, Table S4, S5, Appendix 4), while temperature effects on *Tetrahymena* were exacerbated over time (Fig 3f, perMANOVA *p*<0.01, Table S6, S7, Appendix 4). Both protists showed changes in optical properties as time went by (Fig 3d, e, perMANOVA *p*<0.01), suggestive of changes in cellular contents.

**Figure 3.**
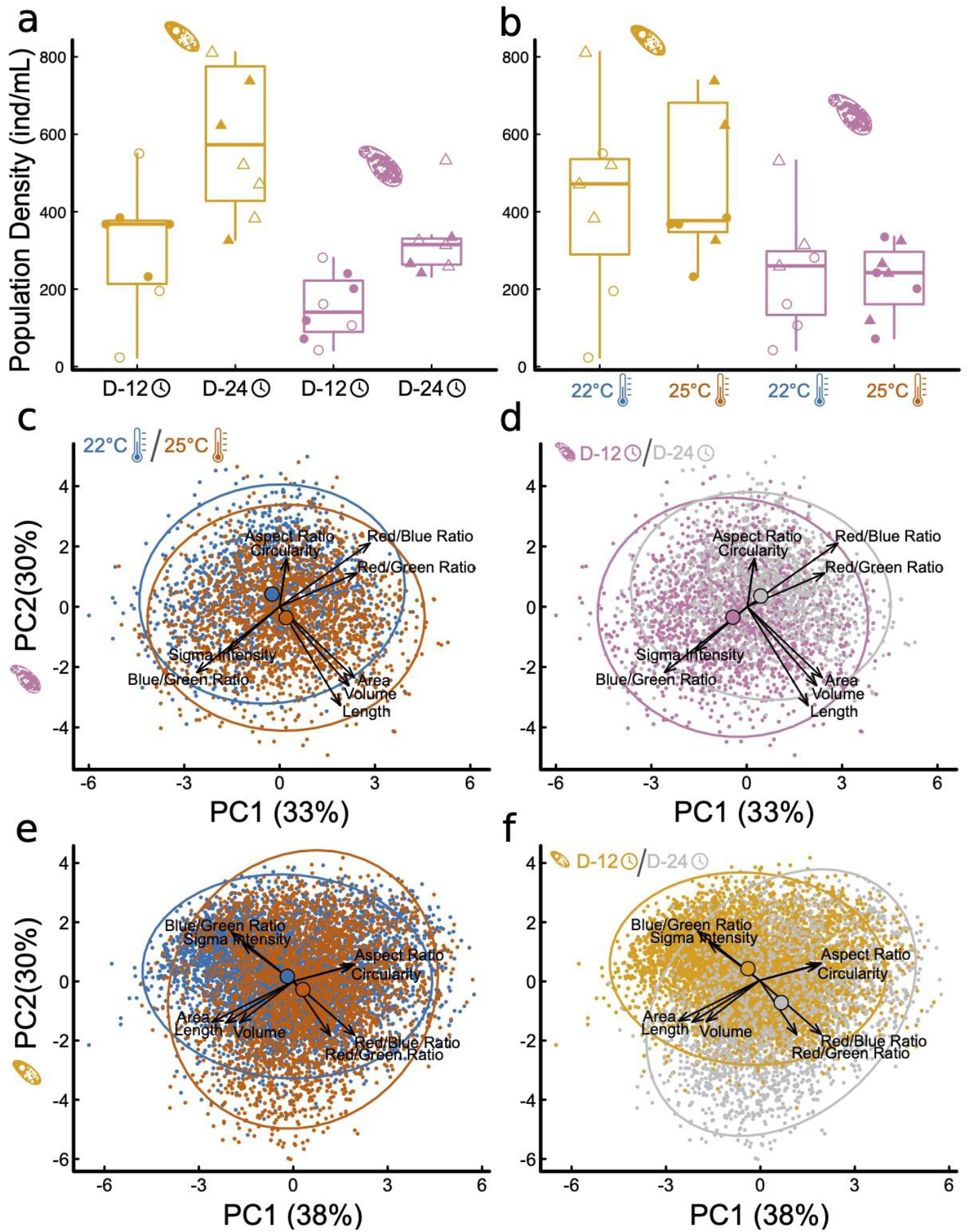
Impact of temperature treatments and incubation time on protist density and morphology. The effect of incubation time (A) and temperature treatment (B) on *Tetrahymena* (yellow) and *Colpidium* (pink) density (cells/mL) in the microcosms; and principal component analysis of multivariate protist phenotypes (C-F): *Colpidium* (C, D) and *Tetrahymena* (E, F), colored by temperature (C, E) and by time (D, F). Vectors represent the principal components loadings of each measured phenotypic characteristic, including: shape, size, optical depth, and hue.

### Density- and Trait Effects of Protists on Taxa-specific Responses

The observed changes in overall bacterial community structure were likely driven by the 113 ASVs that exhibited significant changes in relative abundance (8% of 1423 total ASVs) to the imposed experimental treatments (Fig 4, Appendix 5). Of these responders, 91 ASVs responded significantly to temperature, with 76.9% positively responding to increased temperature and 23.1% showing decreased relative abundance with elevated treatment (Fig 4a). The ASVs flourishing under elevated temperature were largely clustered into several phyla: Verrucomicrobia (12 ASVs), Proteobacteria (10 ASVs), the basal Patescibacteria (9 ASVs), Bacteroidota (7 ASVs) and Spirochaetota (7 ASVs); while the other 25 ‘warm responders’ were spread across ten additional bacterial phyla. In contrast, ASVs thriving under ambient temperatures (21 ASVs) were distributed across the entire bacterial phylogeny (Fig 4a).

**Figure 4:**
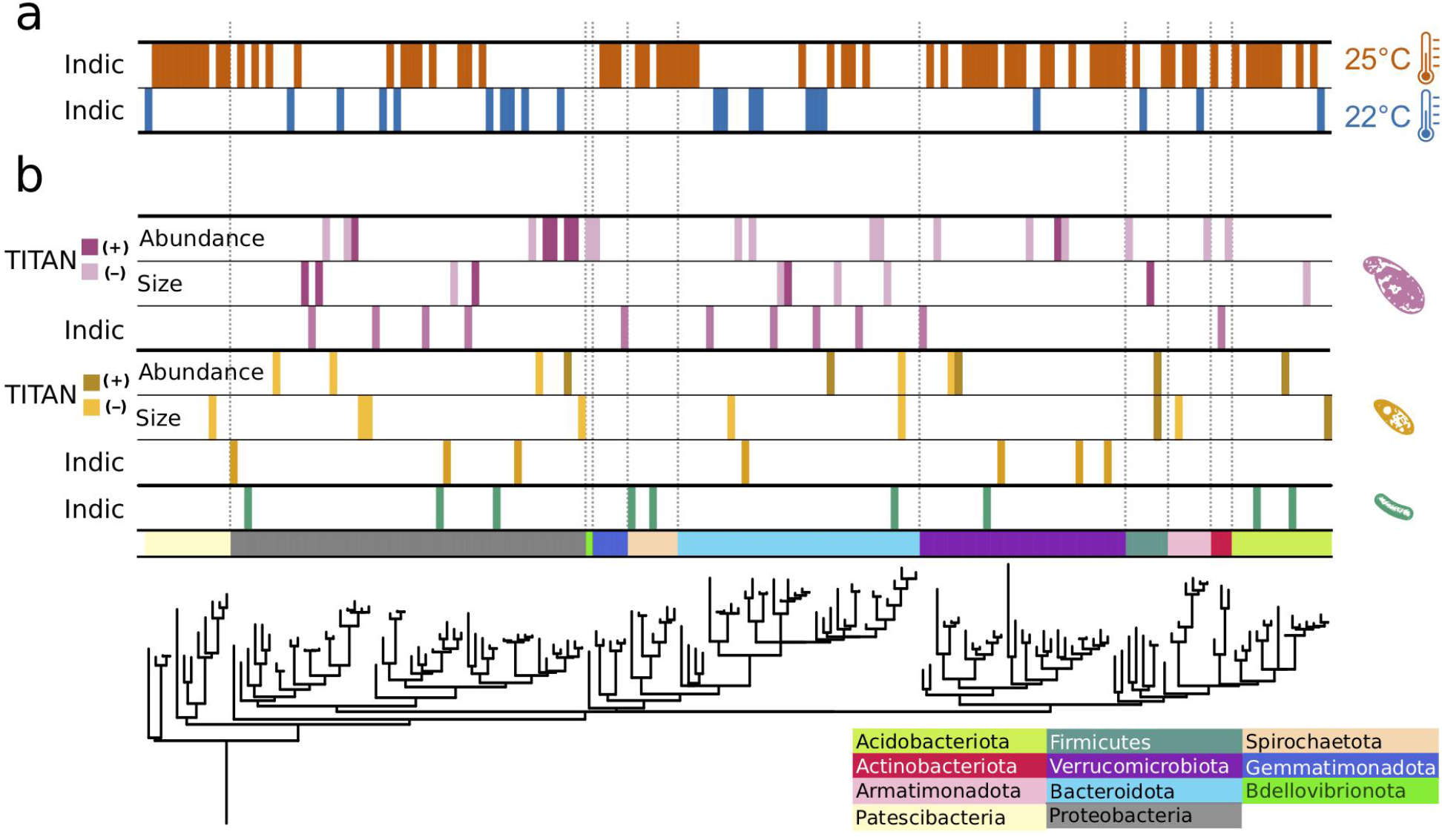
Phylogenetic distribution of treatment impacts on individual bacterial ASVs. Responder ASVs showing consistent change in relative abundance, to temperature (a) and protist (b) treatments. Data strips labeled, “Indic”, represent ASVs positively correlated with a particular temperature treatment level, generated from multi-level pattern analysis output; data strips labeled, “TITAN”, show responding ASVs to changes in specific protist species’ traits, analyzed with direct gradient analysis.

Of the 113 responding ASVs, 23.9% (27 ASVs) exhibited significant shifts in relative abundance to the presence, or absence, of the predator protists (Fig 4b). The presence of *Colpidium* resulted in more responding ASVs (11 ASVs) compared to *Tetrahymena* (7 ASVs) or the no protist treatment (9 ASVs). Responders were distributed throughout the bacterial domain. With TITAN analysis, we also identified 47 ASVs as indicators to gradients of protist cell density and body size (Fig 4b). Seven indicators were negatively associated with *Tetrahymena* body size, while two ASVs positively responded to increased cell size. The density of *Tetrahymena* density corresponded to five positive and five negative responders, distributed across five phyla. Two ASVs responded consistently to cell density and size: a negative indicator ASV_45, identified as *Paenibacillus spp.* (Chitinophagaceae), and a positive responder, ASV_269 in Selenomonadaceae. *Colpidium* cell density and body size resulted in 82% more responding ASVs than to *Tetrahymena* traits. Most of the responders were impacted by *Colpidium* density (21 ASVs), of which the bulk (15 ASVs) responded negatively to more *Colpidium* cells. No ASVs responded to both Colpidium density and cell size, but one responder (ASV 78), in the genus *Afipia*, positively responded to the density of both predators (Fig 4, Appendix 5). ASVs from Proteobacteria, Bacteroidota and Verrucomocrobiota seemed to respond to both protists, while ASVs from Gemmatimonadota and Actinobacteria only responded to Colpidium (Fig 4), thus suggesting some level of specificity to predation by protists, but also to protist species.

## DISCUSSION

Microbes strongly influence the global carbon cycle through respiration and assimilation of both labile and recalcitrant forms of carbon (Jackson et al., 2017). Understanding how changes in environmental temperature may influence microbial community function in general, and respiration rates, in particular, is crucial to hone our ability to forecast future warming trends (REFS). Our study shows how temperature determines both community structure and function in a temperate pond microbial community (Fig 1). We also show how predation by one of the most important consumers of microorganisms worldwide, protist ciliates (Gao et al., 2019; Oliverio et al., 2020), mediate temperature effects on function (Fig 2, 3-4), through Phylum-specific bacterial responses to protist density and size (Fig 4).

Our results show that temperature directly influences community function, owing, in part, to shifts in community structure over time and across temperatures (Fig 2). However, the effects of temperature on function were reversed over time (Fig 2c), which is consistent with other studies showing that while total microbial respiration increases with temperature at first, that effect is short lived or even fully reversed as time elapses (Bradford, 2013; Crowther & Bradford, 2013; Ye et al., 2020; Yergeau et al., 2012). To explain these changes, previous studies have invoked shifts in carbon use efficiency (Frey et al., 2013). While we cannot rule out the possibility of decreased availability of labile carbon within our microcosms, we have attempted to control for that by adding wheat seeds, which provided a slow-release of labile carbon to the incubated communities over time (Altermatt et al., 2015). On the other hand, both temperature and time led to large shifts and increased variability in community structure (Fig 2d), with the warmer temperature leading to a higher relative abundance of some microbial taxa over others (Fig 4), suggesting some degree of phylogenetically conserved responses (Isobe et al., 2020; Martiny et al., 2013). These results thus suggest a possible causal relationship between changes in community structure and function, as proposed by others (Hall et al., 2018), and despite such changes being rarely observed (Fang et al., 2020; Graham et al., 2016; Rocca et al., 2015). While changes in carbon use efficiency have been recently accounted for in state-of-the-art forecasting models (Allison et al., 2010; Sinsabaugh et al., 2013), changes in microbial community structure have not (Sulman et al., 2018). Our results therefore emphasize the need to take such changes into account to improve our predictive models of climate change.

Secondly, we show that predation by larger protists, like *Colpidium sp.*, mediates the impacts of temperature on microbial community function (Fig 2a, b). Indeed, protist predation resulted in changes in total biomass (Fig 2a), microbial respiration that interacted with temperature (Fig 2b) Predation by the larger *Colpidium sp.* actually led to a reversal of the temperature effect on respiration rate (Fig 2b). Interestingly, a recent paper showed that soil decomposition rates increased in the presence of the protist *Physarum polycephalum* at low temperatures, but that effect disappeared at a warmer temperature (Geisen et al., 2021). Our results confirm –and extend– those of Geisen and collaborators to a different protist system, and to a freshwater microbial community, thus suggesting this might be a more general pattern rather than an idiosyncratic effect. If further confirmed, this may also indicate that predation by protists could reduce total microbial respiration in warmer climates, thus representing a poorly understood but potentially important biotic control on the global carbon cycle that ultimately sets the pace of future warming.

Thirdly, protist effects on total microbial biomass (Fig 2a) and function (Fig 2b) were, as hypothesized, size-dependent (Fig 3a,b, Fig 4b). Further analysis also revealed that these effects on microbial communities were likely due to individual bacterial taxa differentially responding to protist density and size (Fig 4b), thus confirming that strong size-dependencies may be at play. Previous studies have shown that protist predation can select for bacterial size (Erktan et al., 2020; Hahn & Höfle, 2001; Pernthaler, 2005), and both negative and positive associations with different protists have also been shown (Oliverio et al., 2020). In addition to the direct predator effects on responding microbial taxa, we cannot rule out indirect effects of the predator presence, due to higher order interactions to explain the abundance shifts of certain responding microbes (Karakoç et al., 2018; Mickalide & Kuehn, 2019). Our results add to this growing literature by mechanistically linking changes in protist density and size to specific bacterial responders (Fig 4b). By showing that cell size and density-dependencies are at play, our experimental results suggest that consumer-resource models that take into account size-dependencies on predation and foraging rates (DeLong et al., 2015; Gauzens et al., 2020; Gilbert et al., 2014) could be used to improve on current Earth System Models once microbial interactions are accounted for.

Lastly, our results showed that the traits of both protist predators also responded to imposed experimental conditions, albeit in different ways. While size is well-known to influence consumer-resource interactions (DeLong et al., 2015; Gauzens et al., 2020; Gilbert et al., 2014), protist size and shape can and often do respond to foraging (Atkinson et al., 2006; DeLong et al., 2014; Gibert et al., 2017; Tan et al., 2021), thus potentially resulting in a feedback between predator and prey phenotypes. Since the microbial communities themselves changed in structure with time and temperature, this study cannot tease apart the direct effects of temperature on protist responses from those mediated by microbial community temperature responses. However understanding how these reciprocal effects may lead to shifts in the function of microbial communities as temperature increases is an exciting avenue for future research.

While our results uncovered interesting patterns about predation influence on the temperature response of microbial communities, there also are shortcomings to our findings that need to be accounted for to fully understand the full scope of these results. For example, the incubation of the microbial communities led to significant departures, in terms of composition, from the original pond microbial communities (Fig S7). One possibility is that our microcosms were artificially awash with nutrients, thus selecting for a microbial composition that would not naturally prevail in more oligotrophic pond conditions. To better understand how common the results shown here may actually be, these abiotic and biotic treatments should also be studied in mesocosms or other semi-natural experimental settings. Our experimental temperature treatments also did not account for diurnal and seasonal temperature fluctuations, nor could they inform us about any broader effects of seasonal changes in temperature. Seasonality, in particular, is likely also shifting as temperatures rise globally (Easterling et al., 2000; Meehl & Tebaldi, 2004; Rahmstorf & Coumou, 2011; Rummukainen, 2012), but whether those effects differ from those of changes in mean temperatures, as shown here, is unknown. Last, the short-term nature of our experimental manipulations necessarily reduce the possible scope of our inference, even though both the nature and magnitude of the effects reported here seem large enough to be of importance beyond our specific set up.

Our results emphasize the dynamic nature of temperature effects on microbial community structure and function as well as how a neglected biological factor (protist predation) influences such responses. We show how protist predation can mediate temperature effects on microbial communities, how such impacts are dependent on the body-size and density of the predator, and how microbial responses to temperature may in turn influence the traits of these microbial consumers. Our study suggests novel and surprising mechanisms that future forecasting models may need to account for to accurately predict biological controls on the global carbon cycle.

## ACKNOWLEDGMENTS

We thank Daniel J. Wieczynski for comments on a previous version of this manuscript. This work was supported by JPG’s Duke University startup funds and a U.S. Department of Energy, Office of Science, Office of Biological and Environmental Research, Genomic Science Program under Award Number DE-SC0020362.

