## Appendix for "Predation by protists influences the temperature response of microbial communities"

Jennifer D. Rocca[1*,2], Andrea Yammine[1], Marie Simonin[3], Jean P. Gibert[1]
[1]Department of Biology, Duke University, Durham, NC, USA
[2]Department of Plant and Microbial Biology, North Carolina State University, Raleigh, NC, USA
[3]University of Angers, Institut Agro, INRAE, IRHS, SFR QUASAV, Angers, France

**INDEX**

**Appendix 1:** Estimation of total respiration rates……………………………………………….. 2-3

**Appendix 2:** Biomass and Respiration Supplemental Results ……………………………….. 4-6

**Appendix 3:** Alpha and Beta-diversity Supplemental Results………………………………… 7-8

**Appendix 4:** Protist traits…………………………………………………………………….. 9-13

**Appendix 5:** Indic Species and TITAN Supplemental Results…… 14 and Table S10 (Separate)

**Appendix 1:** **Estimation of total respiration rates**

Respiration rates were measured as the slope (change over time) of the O2 concentration. Figs S1 and S2 shows the raw data for all jars at D-12 (Fig S1) and D-24 (Fig S2) and the linear regressions from which the slope (respiration rate in micro mol O2.L-1.min-1) was extracted.

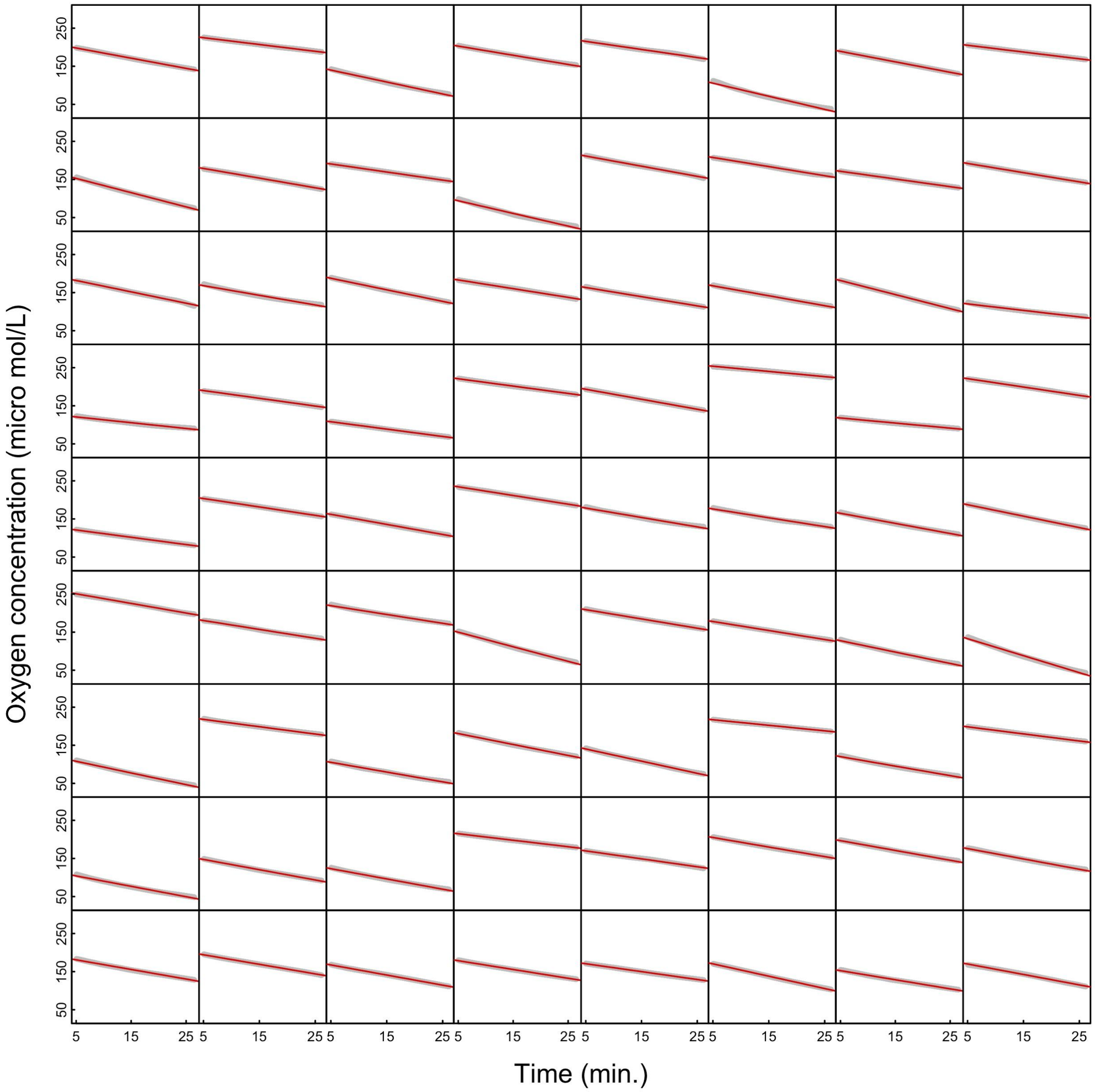

**Fig S1:** Raw data (grey) of all respirometry runs for D-12. Red line represents the regression from which respiration rate (O2 consumed per unit time per L) was extracted.

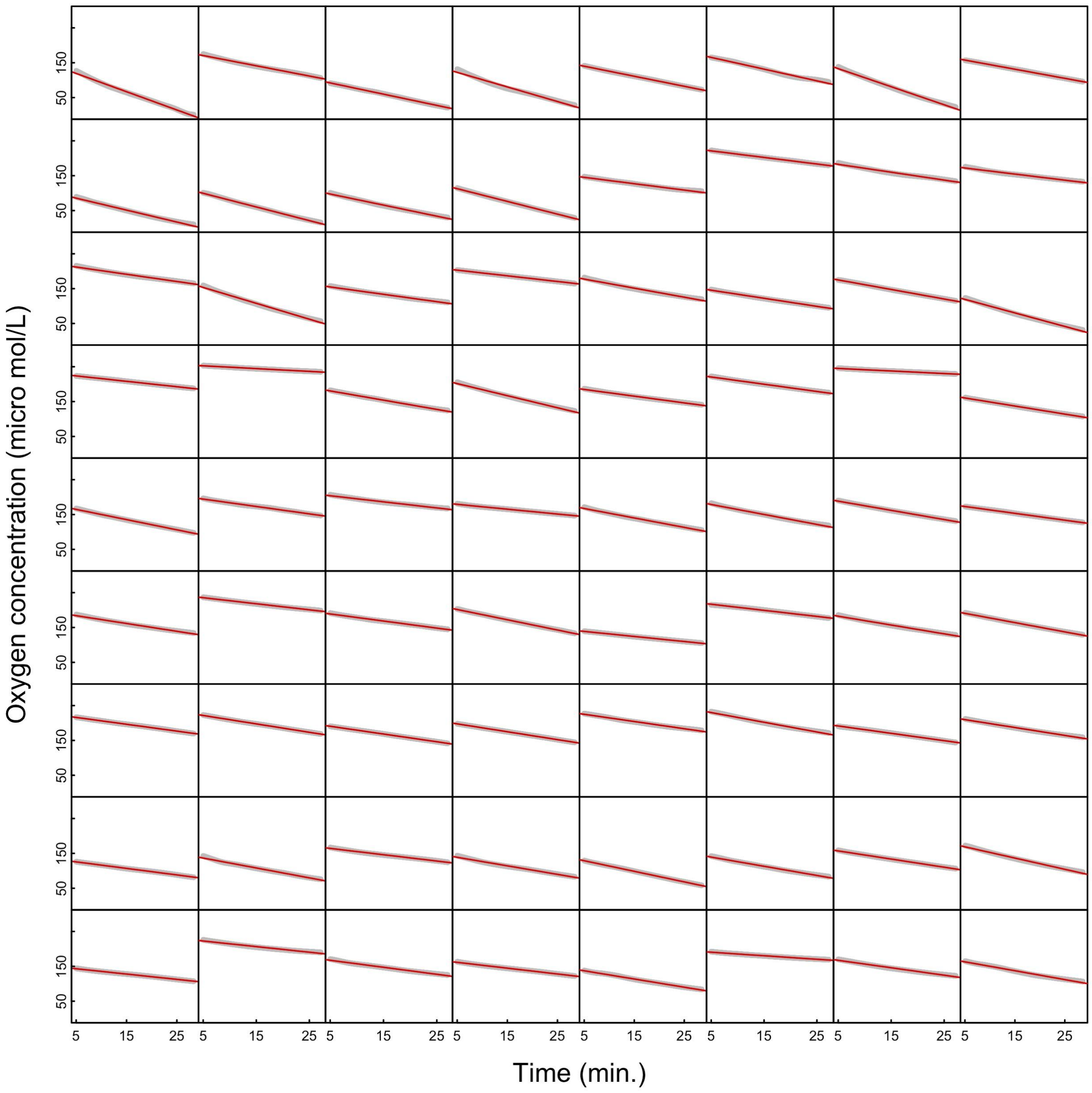

**Fig S2:** Raw data (grey) of all respirometry runs for D-24. Red line represents the regression from which respiration rate (O2 consumed per unit time per L) was extracted.

### <u>Appendix 2:</u> Biomass and Respiration Supplemental Results

*Biomass (as OD600)*

To understand the joint effects of Temperature, Time, and Protists on biomass (OD600), we did exploratory data analysis using the R package "MuMIn". In doing so, we fitted all possible models containing OD600 as the response variable, and all combinations of Temperature, Time, Protist additions (No protists, *Tetrahymena*, *Colpidium*), and their interactions, as explanatory variables. This preliminary data exploration suggested that Protist, Temperature, and Time, were the most important predictors of OD600 (in that order), with the interaction between Temperature and Time coming in fourth (Fig S3, Table S1). Effects reported in the main text were obtained from the best-ranked model.

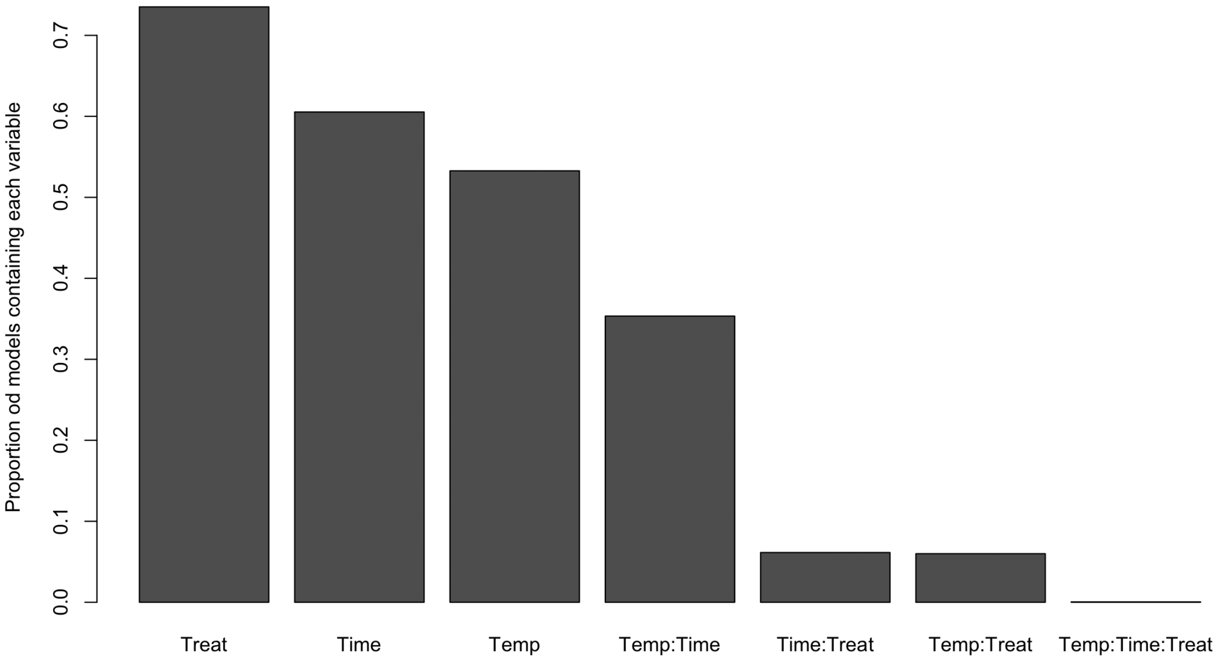

**Fig S3:** MuMIn ranks variables by how often they show up in models with good AICc support. Variables that show up in models with large AICc weight, are considered to be more important than those that don't. This figure shows how Protists (here "Treat"), Time, Temperature, and Time*Temp appear more often in models with low AICc (high AICc weight) than other variables.

**Table S1:** Table of the best models containing the variables Time, Temperature and Protist as well as all possible interactions ranked by AICc (output from “MuMIn” package). It can be seen that the best model accounts for Time, Temperature, Protists, and the interaction between Temperature and Time. However, this model cannot be distinguished using AICc from one that only accounts for Protist treatments. Suggesting all these factors play an important role in determining OD600 (community biomass). However, variance in OD600 is large as well, which is why the null model ranks well.

```
Global model call: lm(formula = Spec ~ Treat * Time * Temp, data = spec_dat, na.action =
"na.fail")
---
Model selection table
```

| | (Int) | Tmp | Tim | Trt | Tmp:Tim | Tmp:Trt | Tim:Trt | Tmp:Tim:Trt | df | logLik | AICc | delta | weight |
|---|---|---|---|---|---|---|---|---|---|---|---|---|---|
| 16 | 1.054 | + | + | + | + | | | | 7 | 45.115 | -75.2 | 0.00 | 0.201 |
| 5 | 1.109 | | | + | | | | | 4 | 41.780 | -75.2 | 0.02 | 0.199 |
| 7 | 1.092 | | + | + | | | | | 5 | 42.317 | -74.1 | 1.12 | 0.114 |
| 1 | 1.060 | | | | | | | | 2 | 38.813 | -73.5 | 1.71 | 0.086 |
| 12 | 1.006 | + | + | | + | | | | 5 | 41.983 | -73.4 | 1.79 | 0.082 |
| 6 | 1.106 | + | | + | | | | | 5 | 41.791 | -73.1 | 2.17 | 0.068 |
| 3 | 1.044 | | + | | | | | | 3 | 39.324 | -72.4 | 2.79 | 0.050 |
| 8 | 1.090 | + | + | + | | | | | 6 | 42.328 | -71.9 | 3.32 | 0.038 |
| 32 | 1.058 | + | + | + | + | + | | | 9 | 45.677 | -71.7 | 3.51 | 0.035 |
| 48 | 1.034 | + | + | + | + | | + | | 9 | 45.552 | -71.5 | 3.76 | 0.031 |
| 2 | 1.058 | + | | | | | | | 3 | 38.824 | -71.4 | 3.79 | 0.030 |
| 39 | 1.073 | | + | + | | | + | | 7 | 42.734 | -70.5 | 4.76 | 0.019 |
| 4 | 1.042 | + | + | | | | | | 4 | 39.336 | -70.3 | 4.91 | 0.017 |
| 22 | 1.111 | + | | + | | + | | | 7 | 42.323 | -69.6 | 5.58 | 0.012 |
| 24 | 1.094 | + | + | + | | + | | | 8 | 42.865 | -68.4 | 6.80 | 0.007 |
| 40 | 1.070 | + | + | + | | | + | | 8 | 42.746 | -68.2 | 7.04 | 0.006 |
| 64 | 1.038 | + | + | + | + | + | + | | 11 | 46.119 | -67.8 | 7.44 | 0.005 |
| 56 | 1.075 | + | + | + | | + | + | | 10 | 43.286 | -64.6 | 10.68 | 0.001 |
| 128 | 1.031 | + | + | + | + | + | + | + | 13 | 46.256 | -63.1 | 12.15 | 0.000 |

Models ranked by AICc(x)

*Respiration rates*

Respiration rates were analyzed like Biomass (OD 600). For each microcosm, respiration rate was quantified twice, once at the temperature treatment at which it had been cultivated, and once at the other temperature treatment to control for possible acute effects of temperature on respiration rates. We therefore tested how all imposed treatments (Temperature, Time and Protists) influenced respiration rates, but also considered the temperature at which the respiration rate was measured. Fig S4 shows that the temperature at which the measurement only minimally influenced the observed respiration rate (if at all), and so it was dropped from all subsequent models.

Interestingly, the effect of the protists was also time dependent, albeit for only one of them. Indeed, microbial communities containing *Colpidium* respired, on average, more over time ($\text{effect}_{\text{Colp}}$= 0.99±0.25SE, $p$<0.01; $\text{effect}_{\text{Colp x Time}}$= 0.94±0.34SE, $p$=0.072; Fig 2b).

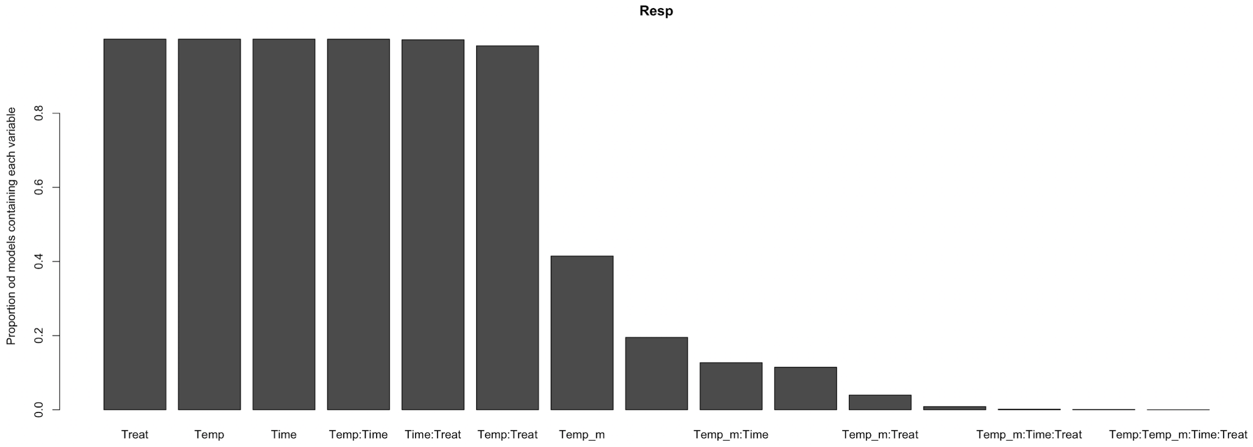

**Fig S4:** Clearly, the most important effects are those of Protists, Temperature of incubation, Time, and then the Temperature and Time interaction, as well as Temperature and Protists and Time and Protists. The temperature at which respiration rates were measured appeared in fewer than 40% of the models, followed by all other interactions.

**<u>Appendix 3:</u> Alpha and Beta-diversity Supplemental Results**

**Table S2:** Effects of time and temperature on alpha diversity. No interactive effects were observed. Alpha diversity estimates calculated on rarefied community data (8620 reads per sample).

| Metric | Effect of Temperature | p-val | Effect of Time | p-val | Effect of Protist | p-val |
|---|---|---|---|---|---|---|
| ASV Richness | +9.1% | 0.01 | +9.2% | 0.01 | NS | 0.624 |
| Faith's phylogenetic | +11.7% | <0.001 | +12.2% | <0.001 | NS | 0.808 |
| Shannon Wiener | +9% | <0.001 | +11% | <0.001 | NS | 0.466 |
| Pielou's Evenness | +7.3% | <0.001 | +9.2% | <0.001 | NS | 0.368 |

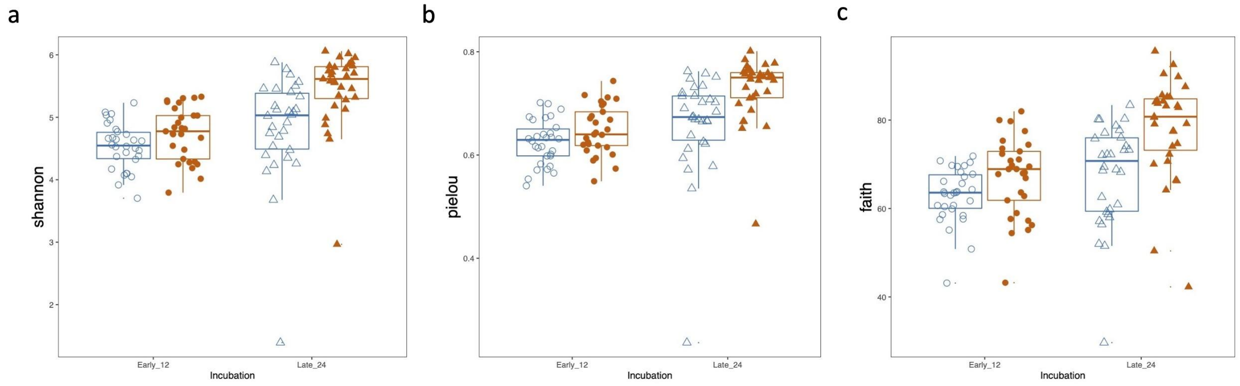

**Figure S5.** We found consistent increases in alpha diversity with time and with elevated temperature across all estimates of alpha diversity: (a) Shannon-Wiener diversity index, (b) Pielou's evenness, and (c) Faith's phylogenetic diversity. Shapes represent incubation time: Day12 - circles, Day 24 - triangles; and temperature marked by point fill: 22$^{o}$C - empty point, 25$^{o}$C - color filled point.

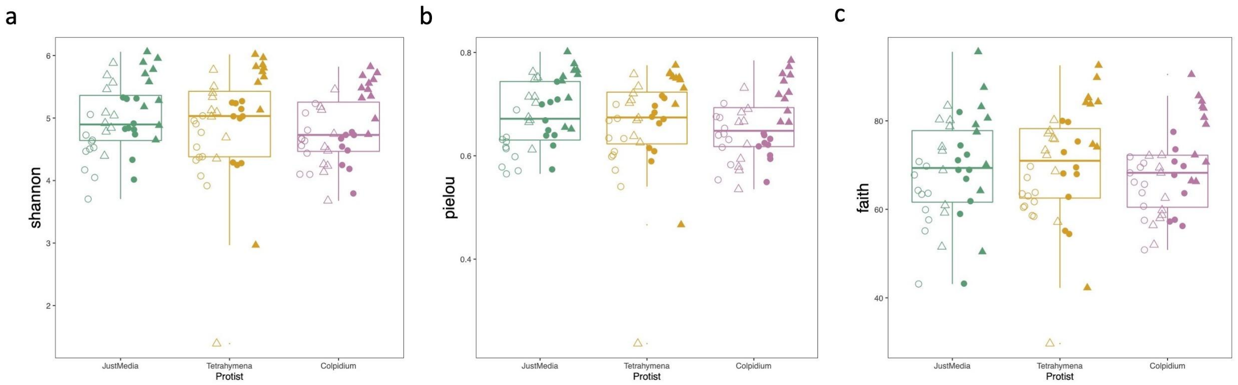

**Figure S6.** Protist presence and species had no impact on any estimate of alpha diversity of the microbial communities: (a) Shannon-Wiener diversity index, (b) Pielou's evenness, and (c) Faith's phylogenetic diversity. Shapes represent incubation time: Day12 - circles, Day 24 - triangles; and temperature marked by point fill: 22$^{o}$C - empty point, 25$^{o}$C - color filled point.

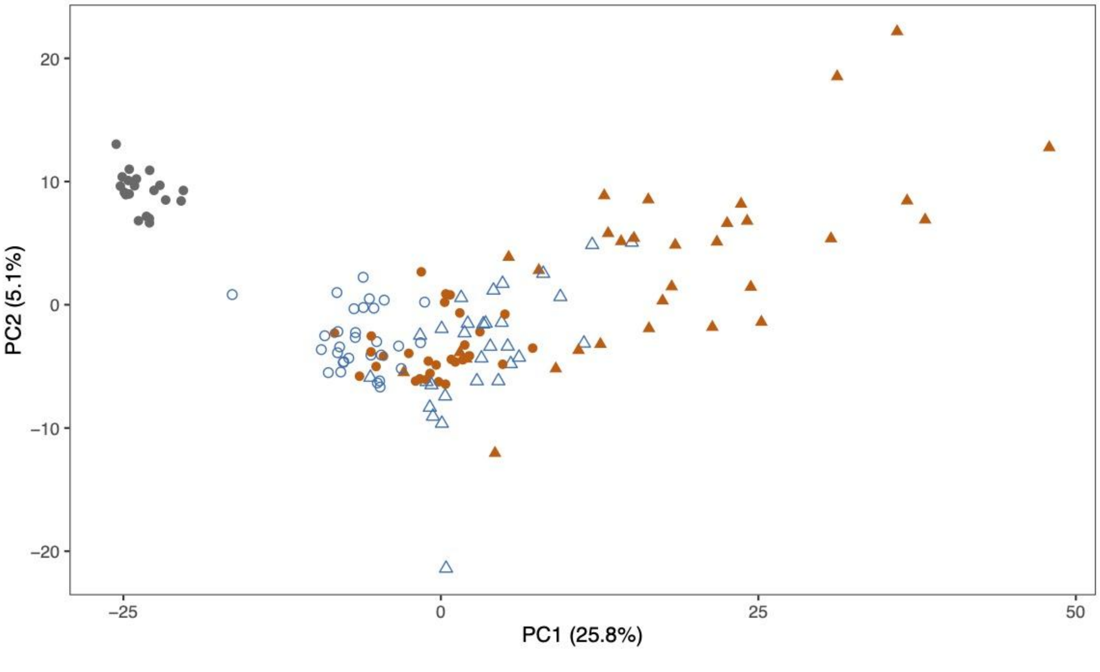

**Figure S7.** Principal component analysis of microbial community structure of the full incubation, including T-0 microbiomes (gray points). Shapes represent incubation time: Day12 - circles, Day 24 - triangles; and temperature marked by point fill: 22$^{o}$C - empty point, 25$^{o}$C - color filled point.

**<u>Appendix 4:</u> Protist trait analyses**

As explained in the main text, we measured multiple protist traits using fluid imaging. To assess how the imposed treatments affected such phenotypes, we used PCA to reduce the dimensionality of the multivariate system. Figs S7 and S8 show how the first three principal components of this multivariate trait space explain over 85% of the total amount of phenotypic variance and where thus retained for analysis.

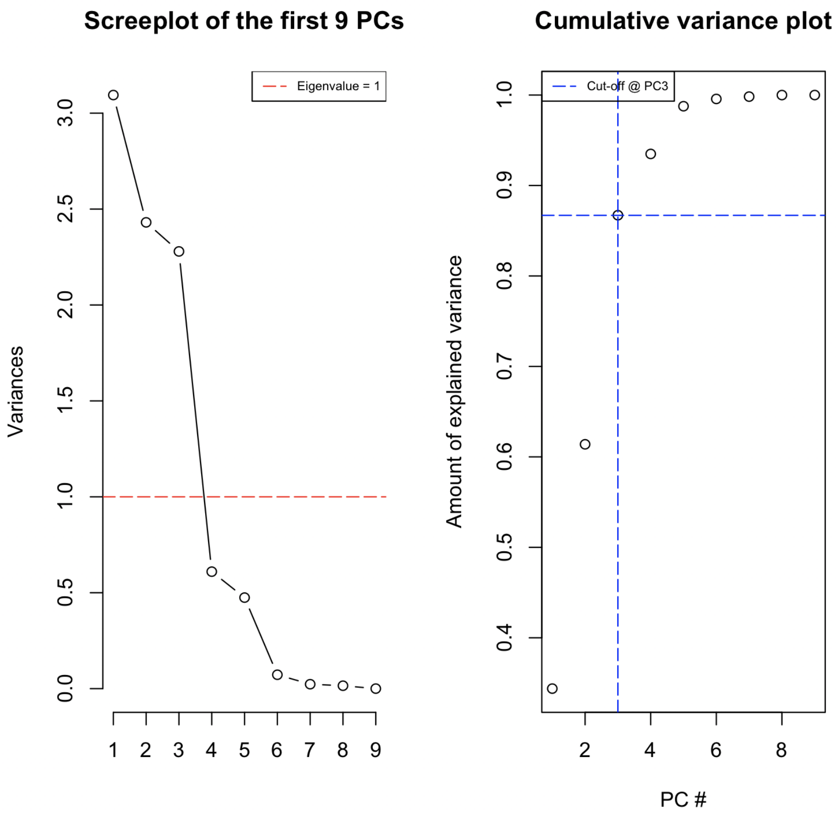

**Fig S8:** Scree plot (left) and cumulative variance plot (right) for *Colpidium* traits. Clearly, the first three principal components explain roughly 85% of the variance, and the first two (shown in the main text), over 60%.

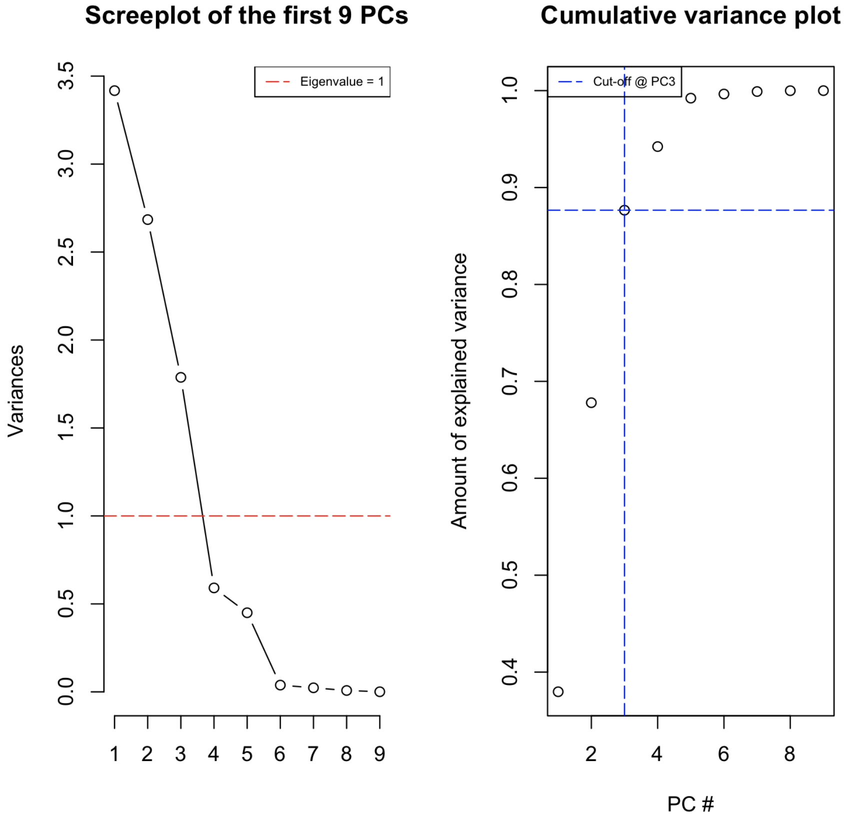

**Fig S9:** Scree plot (left) and cumulative variance plot (right) for *Tetrahymena* traits. Clearly, the first three principal components explain over 85% of the variance, and the first two (shown in the main text), over 65%.

Principal component standard deviation, proportion of total variance explained, and the cumulative proportion of the variance explained are reported for both protists species in Table S3.

**Table S3:** Principal component analysis results for both protist species.

| | PC1 | PC2 | PC3 | PC4 | PC5 | PC6 | PC7 | PC8 | PC9 |
|---|---|---|---|---|---|---|---|---|---|
| <u>*Colpidium*</u> | | | | | | | | | |
| **Stand Dev** | 1.76 | 1.56 | 1.51 | 0.78 | 0.69 | 0.27 | 0.15 | 0.12 | 0.01 |
| **Prop Var Expl** | 0.38 | 0.29 | 0.20 | 0.07 | 0.05 | 0.002 | 0.004 | 0.001 | 0.00 |
| **Cum Var** | 0.38 | 0.68 | 0.88 | 0.94 | 0.99 | 0.99 | 0.99 | 0.99 | 1.00 |
| <u>*Tetrahymena*</u> | | | | | | | | | |
| **Stand Dev** | 1.76 | 1.56 | 1.51 | 0.78 | 0.69 | 0.27 | 0.15 | 0.12 | 0.01 |
| **Prop Var Expl** | 0.34 | 0.27 | 0.25 | 0.07 | 0.05 | 0.008 | 0.003 | 0.002 | 0.00 |
| **Cum Var** | 0.34 | 0.61 | 0.87 | 0.93 | 0.99 | 0.99 | 0.99 | 0.99 | 1.00 |

*perMANOVA*

To assess whether the traits of the protists responded to the imposed temperature (and time) treatments, we used a permutational Multivariate Analysis of Variance (perMANOVA) using R package *vegan*. For both species, we used Time, Temperature, and their interaction, as explanatory variables for the first two principal components. In all cases, temperature, time and their interaction had both additive independent effects and interactive effects (Tables S4-S7).

**Table S4:** perMANOVA table for Colpidium traits PC1

| | Df | SumsOf Sqs | MeanSqs | F.Model | $R^2$ | P-val |
|---|---|---|---|---|---|---|
| Time | 1 | 585.5 | 585.08 | 206.59 | 0.06 | <0.01 |
| Temperature | 1 | 140.3 | 140.34 | 49.55 | 0.015 | <0.01 |
| Time*Temp | 1 | 82.4 | 82.41 | 29.097 | 0.009 | <0.01 |
| Residuals | 3045 | 8623.3 | 2.83 | ——— | 0.91 | ——— |
| Total | 3048 | 9431.6 | ——— | ——— | 1.00 | ——— |

**Table S5:** perMANOVA table for Colpidium traits PC2

| | Df | SumsOf Sqs | MeanSqs | F.Model | $R^2$ | P-val |
|---|---|---|---|---|---|---|
| Time | 1 | 379.1 | 379.14 | 179.03 | 0.05 | <0.01 |
| Temperature | 1 | 476.4 | 476.38 | 224.95 | 0.06 | <0.01 |
| Time*Temp | 1 | 104.8 | 104.79 | 49.48 | 0.01 | <0.01 |
| Residuals | 3045 | 6448.6 | 2.12 | ——— | 0.87 | ——— |
| Total | 3048 | 7408.9 | ——— | ——— | 1.00 | ——— |

**Table S6:** perMANOVA table for Tetrahymena traits PC1

| | Df | SumsOf Sqs | MeanSqs | F.Model | $R^2$ | P-val |
|---|---|---|---|---|---|---|
| Time | 1 | 2098.3 | 2098.28 | 693.93 | 0.08 | <0.01 |
| Temperature | 1 | 79.9 | 79.87 | 26.41 | 0.003 | <0.01 |
| Time*Temp | 1 | 992.1 | 992.09 | 328.10 | 0.036 | <0.01 |
| Residuals | 8025 | 24265.5 | 3.02 | ——— | 0.88 | ——— |
| Total | 8028 | 27435.7 | ——— | ——— | 1.00 | ——— |

**Table S7:** perMANOVA table for Tetrahymena traits PC2

| | Df | SumsOf Sqs | MeanSqs | F.Model | $R^2$ | P-val |
|---|---|---|---|---|---|---|
| Time | 1 | 2478.1 | 2478.08 | 1054.98 | 0.11 | <0.01 |
| Temperature | 1 | 28.3 | 28.32 | 12.06 | 0.001 | <0.01 |
| Time*Temp | 1 | 193.7 | 193.66 | 82.45 | 0.009 | <0.01 |
| Residuals | 8025 | 18850.1 | 2.35 | ——— | 0.87 | ——— |
| Total | 8028 | 21550.2 | ——— | ——— | 1.00 | ——— |

**Appendix 5: Indic Species and TITAN Supplemental Results**

With indicator species analysis on the categorical treatments as well as TITAN on gradients of the protist traits, we identified 157 microbial taxa (ASVs) responding significantly to the incubation. Changes in relative abundance were identified from the variance-stabilizing transform of the community sequencing dataset, to account for differences in sequencing depth as well as to treat the data compositionally. Among the indicator species analysis, we identified just five ASVs that responded positively to more than one treatment level: ASV499 (vandinBE97) increased in 25$^{o}$C and no protist; ASV535 (*Pedobacter*) increased in 22$^{o}$C and *Colpidium* presence; ASV563 (*Ferrovibrio*) increased in 25$^{o}$C and *Colpidium*; ASV738 (Bryobacter) increased in 25$^{o}$C and Tetrahymena; ASV833 (Pedosphaeraceae) increased in 25$^{o}$C and Tetrahymena. Finally, no ASVs responded both to the presence of a protist predator and to that protist's corresponding cell size or density.

**Table S10. Taxonomic assignment for responding ASVs.** Silva 138-assigned taxonomy for the responding ASVs, and corresponding response directionality. *See separate spreadsheet file, containing the ASV names, taxonomic assignments (based on Silva 138) and response categories.*
